# Phosphoglycerol-type wall- and lipoteichoic acids are enantiomeric polymers differentially cleaved by the stereospecific glycerophosphodiesterase GlpQ

**DOI:** 10.1101/2020.01.08.899310

**Authors:** Axel Walter, Sandra Unsleber, Jeanine Rismondo, Ana Maria Jorge, Andreas Peschel, Angelika Gründling, Christoph Mayer

## Abstract

The cell envelope of Gram-positive bacteria generally comprises two types of polyanionic polymers, either linked to peptidoglycan, wall teichoic acids (WTA), or to membrane glycolipids, lipoteichoic acids (LTA). In some bacteria, including *Bacillus subtilis* strain 168, WTA and LTA both are glycerolphosphate polymers, yet are synthesized by different pathways and have distinct, although not entirely understood morphogenetic functions during cell elongation and division. We show here that the exo-lytic *sn*-glycerol-3-phosphodiesterase GlpQ can discriminate between *B. subtilis* WTA and LTA polymers. GlpQ completely degrades WTA, lacking modifications at the glycerol residues, by sequentially removing glycerolphosphates from the free end of the polymer up to the peptidoglycan linker. In contrast, GlpQ is unable to cleave unmodified LTA. LTA can only be hydrolyzed by GlpQ when the polymer is partially pre-cleaved, thereby allowing GlpQ to get access to the end of the polymer that is usually protected by a connection to the lipid anchor. This indicates that WTA and LTA are enantiomeric polymers: WTA is made of *sn*-glycerol-3-phosphate and LTA is made of *sn*-glycerol-1-phosphate. Differences in stereochemistry between WTA and LTA were assumed based on differences in biosynthesis precursors and chemical degradation products, but so far had not been demonstrated directly by differential, enantioselective cleavage of isolated polymers. The discriminative stereochemistry impacts the dissimilar physiological and immunogenic properties of WTA and LTA and enables independent degradation of the polymers, while appearing in the same location; e.g. under phosphate limitation, *B. subtilis* 168 specifically hydrolyzes WTA and synthesizes phosphate-free teichuronic acids in exchange.

## Introduction

Bacteria are covered by a complex multilayered cell envelope positioned external to the cell membrane, which protects the susceptible protoplast from detrimental effects of the environment and the cells from lysis (1). Differences within the composition of the cell envelope classifies the two major groups of bacteria, Gram-negatives and Gram-positives. Gram-negative bacteria are encased in a thin peptidoglycan (PGN) layer that is covered by an external outer membrane (OM), carrying negatively charged lipopolysaccharide (LPS) in the outer leaflet. In contrast, Gram-positive bacteria lack an OM, but possess a thick PGN layer that is interweaved by polyanionic glycopolymers, the teichoic acids, which were discovered by Baddiley and coworkers 60 years ago (2–4). Teichoic acids can be very variable in composition and structure, although they mostly feature glycerol-phosphate, ribitol-phosphate, or sugar phosphate repeating units connected through phosphodiester bonds (5–8). These phosphodiester-polymers are either covalently bound to the PGN and called wall teichoic acids (WTA) or linked to membrane glycolipids, anchored in the cell membrane and named lipoteichoic acids (LTA) (4,9,10). WTA are characteristic constituents of the Gram-positive cell walls (PGN-WTA complex), comprising chains of 30 to 50 polyol-phosphate repeats, connected via phosphodiesters and anchored via a linker disaccharide *(N*-acetylmannosamine-β-1,4-*N*-acetylglucosamine; ManNAc-β-1,4-GlcNAc) to about every ninth *N*-acetylmuramic acid (MurNAc) residue of the PGN (9). They make up about half of the cell wall dry weight (11,12) and are responsible for the generally high phosphate content of Gram-positive cell walls (4,13). It was shown that WTA can serve as phosphate storage, allowing *B. subtilis* to continue growth under phosphate-depleted conditions (13–15). Under these conditions, teichoic acids are exchanged with phosphate-free teichuronic acids to cope with this stress, which is an adaptation process known as the “teichoic acid-to-teichuronic acid switch”. LTA is more widespread in bacteria than WTA and the composition is less dependent on growth conditions (16). Commonly, LTA contain polyol-phosphate chains (Type I LTA) that are anchored in the cytoplasmic membrane via glycolipids; in the case of *B. subtilis*, a gentibiosyl disaccharide (glucose-β-1,6-glucose) glycosidically bound to diacylglycerol (10). Although differences in the chemical composition, route of biosynthesis, as well as roles in cell growth and morphogenesis have been identified between WTA and LTA, their physiological functions are still insufficiently understood (4,17–19). Inactivation of both LTA and WTA is lethal in *B. subtilis*, indicating partially redundant functions, nevertheless comparison of the individual mutants suggested that they have distinct roles during cell elongation (WTA) and division (LTA) (18). Further proposed functions of teichoic acids include control of cell wall targeting enzymes during envelope homeostasis and divalent cation binding (3,20), interaction with host and bacteriophage receptors (18,21), as well as pathogenicity (22–24). Recently, LTA has been suggested to functionally resembling the osmoregulated periplasmic glycans of Gram-negative bacteria (10,25,26).

In some Gram-positive bacteria, including *B. subtilis* 168 as well as in *S. epidermidis* and *S. lugdunenis*, both LTA and WTA are glycerophosphate polymers (17,18,27,28). Nevertheless, they are synthesized by distinct routes (9,10,29). Figure 1 summarizes the biosynthesis pathways of WTA and LTA of *B. subtilis* 168. WTA is synthesized from CDP-glycerol in the cytoplasm and the polymer is then flipped outwards (Fig. 1A). In contrast, LTA is synthesized from the precursor phosphatidylglycerol (PG), generated via diacylglycerol(DAG)-CDP and PG-phosphate (PGP). The PG precursor is subsequently translocated and then polymerized on the outside of the cell (Fig. 1B). An important finding has been that the glycerophosphate in the precursors of WTA and LTA have different stereochemistry (17). CDP-glycerol has a *sn-*3-configuration, and in contrast to this, the free glycerol phosphate of PG has a *sn-*1-configuration. The prochirality of glycerol leads to two 3-phosphate products: according to convention (stereochemical numbering: *sn-*nomenclature), L-glycerol is the configuration that determines the numbering of *sn*-glycerol phosphates (Fig. 1C). The use of different precursors and the compartmentalization of WTA and LTA synthesis allows to differentially regulate the production of the two polymers, which is important in order to fulfil particular roles in cell envelope integrity and distinct morphogenetic functions during cell cycle and growth phases (18). In contrast, how WTA and LTA execute their distinct physiological functions is not obvious, as they are both present in the same cell envelope compartment.

**Figure 1.**
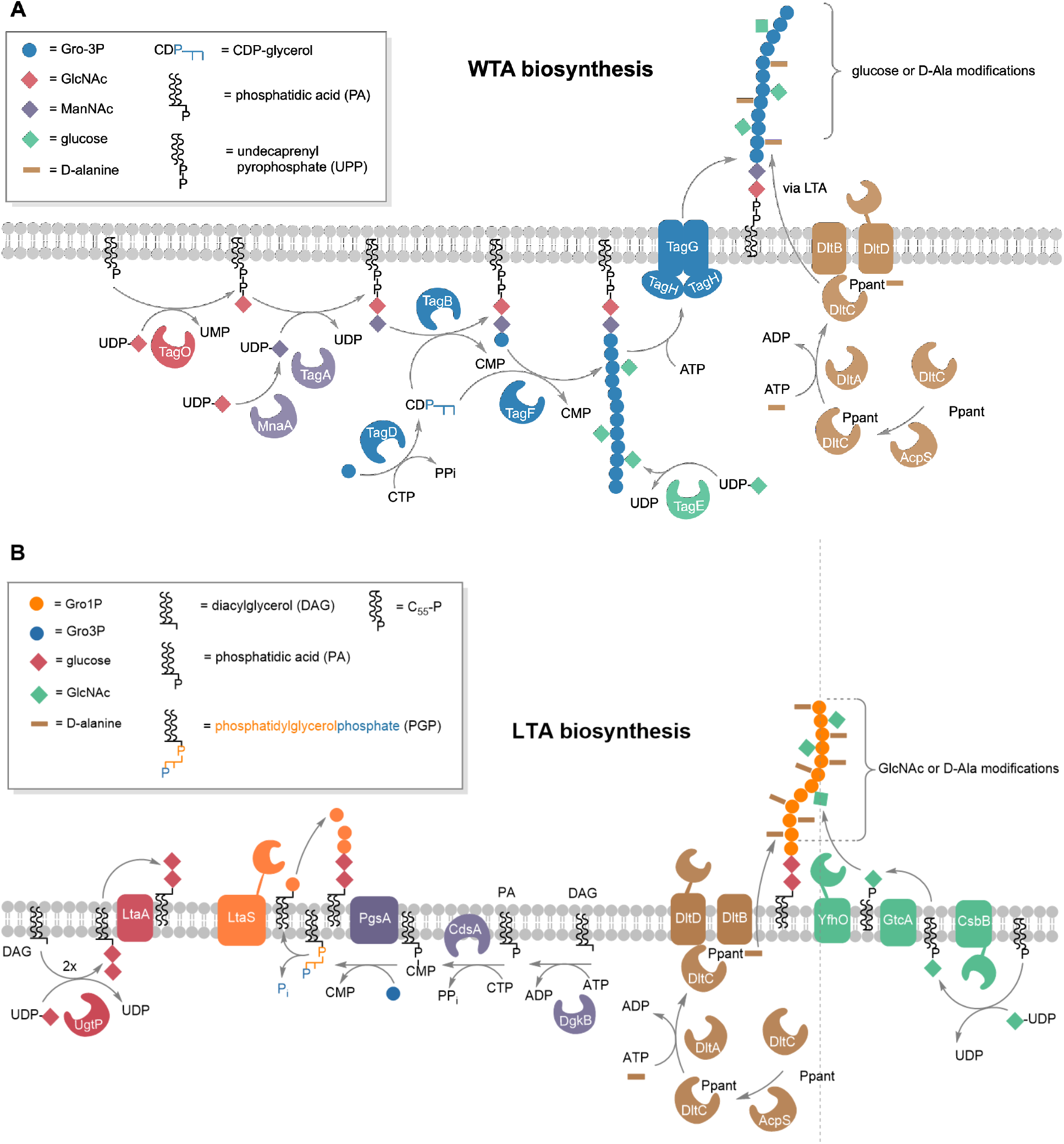
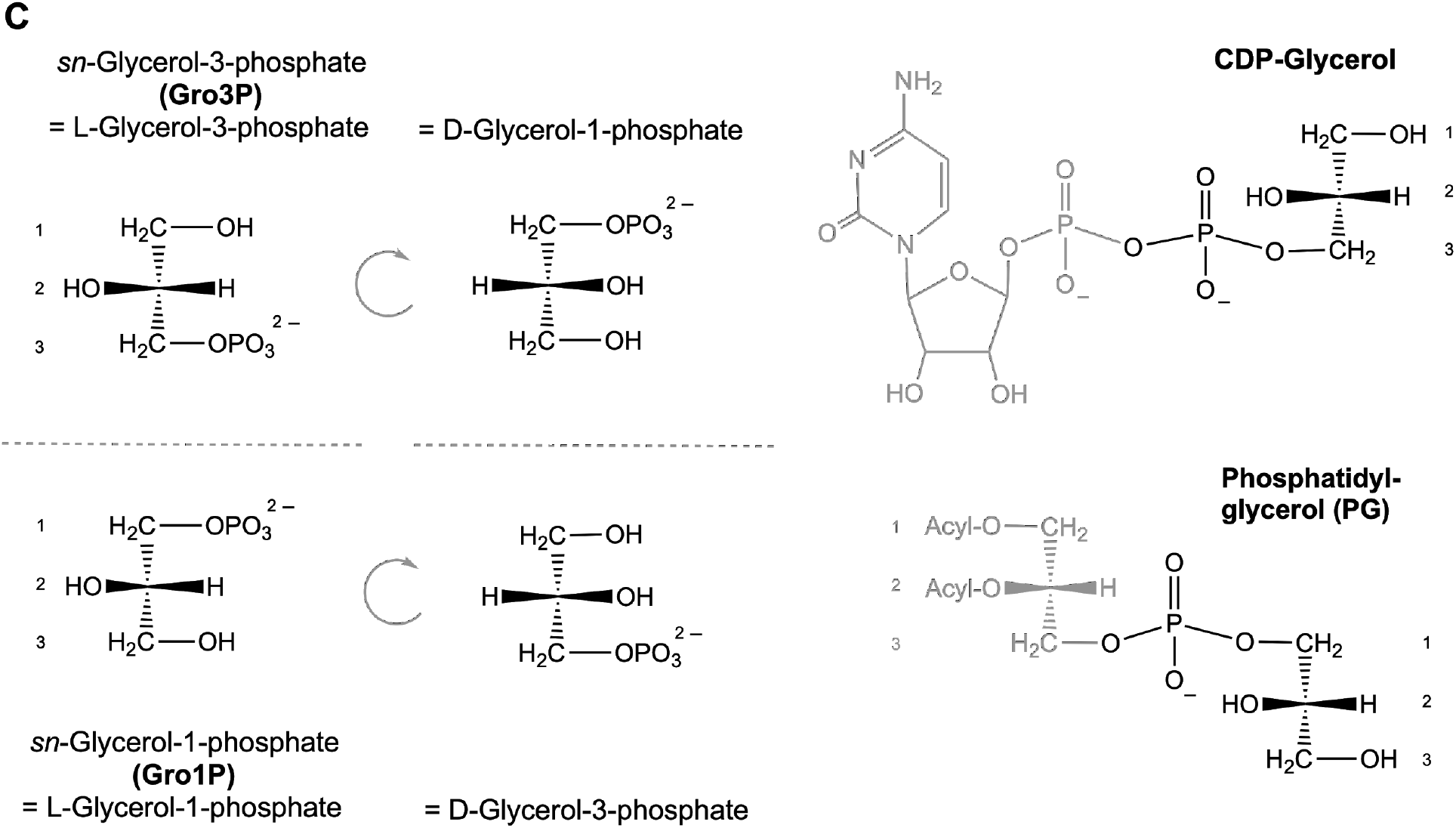
Comparison of the biosynthesis pathways of WTA and LTA and the stereochemistry of glycerol phosphates and teichoic acid precursors. ***A***, Overview of WTA biosynthesis. TagO initiates WTA biosynthesis by transferring GlcNAc (red diamond) from UDP-GlcNAc onto the lipid carrier undecaprenyl phosphate while releasing UMP. MnaA converts UDP-GlcNAc to UDP-ManNAc, which in turn is used to transfer ManNAc (purple diamond) to the TagO product, thereby forming the WTA linker disaccharide bound to undecaprenyl pyrophosphate (UPP). The TagB protein catalyzes a priming step of WTA synthesis in *B. subtilis*, thereby completing the linkage unit (GlcNAc-ManNAc-Gro3P): a single Gro3P (blue circle) is added from CDP-glycerol (which is generated by TagD from Gro3P and CTP) to the membrane-anchored linker disaccharide, while releasing CMP. TagF further elongates the WTA chain polymer by repeatedly transferring Gro3P (blue circles) from CDP-glycerol. The Gro3P units of the WTA polymer get partially glycosylated in the cytoplasm. The enzyme TagE utilizes UDP-glucose to attach glucose (green diamonds) onto the C2 hydroxyl group of Gro3P of the chain polymer. The degree of glycosylation strongly depends on growth conditions and growth phase. Export of the WTA polymer through the cell membrane is achieved by the ABC-transporter TagGH. Eventually, the DltABCD system attaches D-alanyl esters to unglycosylated parts of WTA, which has been reported to occur indirectly via LTA (34). DltA transfers D-alanine in a ATP-dependent two-step reaction to DltC, which has been modified with 4’-phosphopantetheine (Ppant) at Ser35 by acyl carrier protein synthase (AcpS) (56). DltB interacts with DltC-Ppant and transfers the D-alanyl onto the C2 hydroxyl group of Gro3P of teichoic acid chains (10,57). ***B***, Overview of LTA biosynthesis. The LTA precursor phosphatidyl glycerolphosphate (PGP) is generated by a series of reactions within the cytoplasm. Synthesis starts by phosphorylation of the lipid carrier diacylglycerol (DAG) yielding DAG-phosphate (= phosphatidic acid; PA). The enzyme CdsA then transfers a CMP moiety from CTP onto PA, yielding DAG-CDP while releasing pyrophosphate (PP_i_). The CMP moiety of the latter gets exchanged with Gro3P catalyzed by PgsA, thereby forming PGP. Notably, phosphatidylglycerol (PG) is formed by releasing the *sn*-3-phosphoryl group from PGP, thereby retaining a Gro1P entity. It is unknown if this reaction is catalyzed by a so far uncharacterized phosphatase, or if rather the LTA synthase LtaS catalyzes this reaction, in order to energize the polymerisation of Gro1P entities, which are added to the lipid-anchored linker disaccharide in the outer leaflet of the plasma membrane by the housekeeping LTA synthase LtaS_BS_. The linker disaccharide (red diamonds) is synthesized by UgtP, adding two glucose from UDP-glucose onto DAG. LtaA flips the DAG-linker across the membrane (58). Besides LtaS_BS_, *B. subtilis* also contains further LTA synthases that catalyse the polymerization of the LTA chains (orange circles): the stress-induced (YfnI) and the sporulation-specific (YqgS) LTA synthase as well as the LTA primase YvgJ, which adds an initial Gro1P to the disaccharide linkage unit (59). LTA polymers may get modified by D-alanylation (brown) and glycosylation (green). As described above alanylation is catalyzed by the Dlt alanylation system: DltA transfers D-alanine to DltC-Ppant and subsequently DltB transfers the D-alanyl onto the C2 of Gro1P (10,57). Glycosylation of LTA is catalyzed by the glycosyltransferase CsbB that adds GlcNAc onto undecaprenyl phosphate (C_55_-P) using UDP-GlcNAc. As this modification occurs outside the cell in *B. subtilis*, C_55_-P-GlcNAc is first flipped across the membrane by the flippase GtcA and then the glycosyltransferase YfhO modifies the C2 of Gro1P with GlcNAc (38,40). ***C***, The precursors of WTA and LTA synthesis carry enantiomeric glycerol phosphates: *sn*-glycerol 3-phosphoryl (CDP-glycerol) and *sn*-glycerol 1-phosphoryl group (phosphatidyl-glycerol; PG), respectively. Glycerol phosphate enantiomers are defined by convention according to stereochemical numbering (sn-nomenclature) as *sn*-glycerol-3-phosphate (Gro3P = L-glycerol-3-phosphate = D-glycerol-1-phosphate) and *sn*-glycerol-1-phosphate (Gro1P = L-glycerol-1-phosphate = D-glycerol-3-phosphate).

Modifications of the polyols by alanylation and glycosylation are important means to alter the physiological properties of WTA and LTA and also affect recognition by the innate immune system (6,30,31). D-alanylation adds positive charge (free amino groups) to the polyol-phosphate polymers, thereby rendering the anionic character and as a consequence the binding properties (10,32). The multienzyme complex DltABCD is responsible for adding D-Ala modifications onto LTA on the outer leaflet of the cell membrane, and indirectly also onto WTA (Fig. 1) (32–35). LTA and WTA can also be α-, or β-glycosylated, which severely increases the stability of these polymers against alkaline hydrolysis (9,10). In *B. subtilis* the enzyme TagE transfers α-glucosyl residues from UDP-glucose onto preformed WTA within the cytoplasm and constitutes the only WTA glycosylating enzyme in this bacterium (Fig. 1A) (36,37). Although WTA glycosylation usually occurs prior to the translocation of the polymer across the cell membrane, it was recently proposed that it may also occur after translocation in *Listeria monocytogenes* (37,38). Alike alanylation, glycosylation of LTA generally occurs, along with synthesis, outside the cell, and membrane associated three component glycosylation systems responsible for LTA glycosylation have been characterized recently in *B. subtilis* and *S. aureus (*CsbB/GtcA/YfhO) as well as in *L. monocytogenes* (GtlA/GtlB) (Fig. 1B) (38–40).

Besides synthesis, also the turnover of WTA and LTA needs to be differentially regulated, which so far has not been explored in much detail. Recently, the exo-acting *sn*-glycero-3-phosphate phosphodiesterase GlpQ, along with an endo-acting phosphodiesterase PhoD, has been implicated in the degradation of WTA during phosphate starvation (41). However, apart from WTA degradation during adaptation to phosphate starvation, turnover of WTA likely occurs also along with the turnover of PGN of the cell wall in *B. subtilis* and other Gram-positive bacteria (42–44). Since strains of *B. subtilis* lacking both WTA and LTA are not viable, the simultaneous degradation of both polymers would be detrimental (18). We thus wondered how differential degradation of WTA and LTA by hydrolases (“teichoicases”) is possible. Previous studies with the glycerophosphodiesterase GlpQ of *B. subtilis* as well as orthologous enzymes from *Escherichia coli* and *S. aureus* (amino acid sequence identities of 29 and 54 %, respectively) have revealed strict stereospecificity for glycerophosphodiesters harbouring *sn*-glycerol-3-phosphoryl groups, e.g. produced by phospholipases from membrane phospholipids (41,45–47). Accordingly, phosphatidylglycerol or lysophosphatidylglycerol, which harbour only free *sn*-glycerol-1-phosphoryl ends are not hydrolyzed by GlpQ and also bis(p-nitrophenyl)-phosphate, a chromogenic substrate for other phosphodiesterases, is not cleaved by GlpQ (45,46). Intriguingly, LTA of *S. aureus* was found to be not a substrate of GlpQ, which however could be due to phosphoglycerol backbone modifications (47). In contrast, the enzyme shows broad substrate specificity with respect to the alcohol moiety and can hydrolyse a variety of different phospholipid head groups, such as glycerophosphocholine, glycerophosphoethanolamine, glycerophosphoglycerol, and bis(glycerophospho)glycerol (41,45,47).

So far, differential cleavage of WTA and LTA polymers by GlpQ has not been examined in detail. In this work, we show that the stereospecific *sn*-glycerol-3P phosphodiesterase GlpQ acts as an exo-lytic hydrolase that sequentially cleaves off *sn*-glycerol-3-phosphate (Gro3P) entities from the exposed end of WTA, however, it is unable to hydrolyse intact LTA. Thereby we provide biochemical evidence that these polymers have opposite stereochemistry: WTA constitute phosphodiester-polymers made of Gro3P and LTA polymers of *sn*-glycerol-1-phosphate (Gro1P). The stereochemical difference likely impacts many of the polymers’ distinct properties, such as interactions with hydrolases and binding of proteins throughout the cell cycle, bacterial growth and differentiation.

## RESULTS AND DISCUSSION

### GlpQ is a stereospecific *sn*-glycerol-3-phosphoryl phosphodiesterase

GlpQ of *B. subtilis* and orthologs from other bacteria have been shown previously to specifically release Gro3P from GPC, glycerophosphoethanolamine, glycerophosphoglycerol, and bis(glycerophospho) glycerol. For the latter two substrates, *K*_M_ and *k*_cat_ values of 1.0 mM and 1275 min^−1^, respectively 1.4 mM and 1517 min^−1^, were determined for *B. subtilis* GlpQ (41,45,47). We confirmed the stereospecificity of recombinantly expressed, *B. subtilis* GlpQ for *sn*-glycero-3-phosphoryl substrates and determined the enzyme’s stability and catalytic optima, using *sn*-glycero-3-phosphocholine (GPC) as substrate (Supporting Information, Fig. S1). Our analysis revealed that GlpQ is rather temperature sensitive. It readily loses stability at temperatures above 30°C, e.g. within 30 min at 37°C more than 50% of its activity was lost. At the same time however, the enzymatic turnover steadily increases with temperature up to an optimum at 55°C and about half maximum activity at 30°C (Supporting Information, Fig. S1B). Furthermore, the enzyme was shown to be stable over a remarkably wide pH range between 2 to 10, but has a very narrow optimum at pH 8.0 (Supporting Information, Fig. S1B). We thus conducted all experiments with the enzyme GlpQ in this study at 30°C and pH 8.0.

Although the detailed mechanism of phosphodiester-cleavage by GlpQ is currently unknown, Ca^2+^ ions were recognized as crucial for catalytic activity (yet they can be substituted with Cd^2+^ and partially with Mn^2+^ and Cu^2+^) (45,48). Accordingly, the catalytic reaction was inhibited with ethylenediaminetetraacetic acid (EDTA). Nevertheless the addition of Ca^2+^ ions was not required when using the recombinant GlpQ that was purified from the cytosolic extracts of *E. coli*. The recently solved crystal structure of the *B. subtilis* GlpQ with Gro3P bound to the active site (structural database identifiers: 5T9B and 5T9C) confirmed the importance of a Ca^2+^ ion for catalysis as well as for the stereospecific coordination of the substrate (41,48). The active site of GlpQ includes a residue (His85) that is located on a small additional, so-called glycerophosphodiester phosphodiesterase domain, which is inserted between the beta-strand and alpha-helix of the second beta/alpha motif of a classical triose phosphate isomerase (TIM)-barrel structure (41,49). Figure 2 depicts the substrate and Ca^2+^ binding sites of GlpQ, located in a deep pocket located on the TIM barrel domain, and rationalizes the strict stereospecificity of the enzyme. The substrate binding cleft can be divided into a hydrophilic side including the active site Ca^2+^ ion and a hydrophobic side consisting of hydrophobic amino acids including phenylalanine and tyrosine (Phe190, Tyr259, Phe279) (Fig. 2). The active site Ca^2+^ ion adopts a pentagonal bipyramidal coordination. It is held in place by glutamic and aspartic acid residues (Glu70, Glu152, Asp72) and is also coordinated by the two hydroxyl groups of Gro3P (Fig. 2). The phosphate as well as the C2 and C3 hydroxyl-group of Gro-3P are drawn towards the active site Ca^2+^ ion, and are moved away from the hydrophobic side of the binding cleft. Coordination of the Ca^2+^ ion by amino acids with charged side chains and the hydroxyl and phosphate groups of the substrate as well as the orientation of the hydrophobic C-H groups of the substrate towards the hydrophobic side of the binding cleft restricts the productive binding to the unsubstituted *sn*-glycero-3-phosphoryl stereoisomer, thus allowing productive binding only of *sn*-glycerol-3-phosphoryl groups. Instead, the C2 hydroxyl group of *sn*-glycerol-1-phosphoryl would face towards the hydrophobic side, making the binding impossible. The hydrophilic side of the binding cleft also coordinates the phosphate group of the substrate involving the basic side chains of His43, Arg44, His85 (Fig. 2). His43 and His85 presumably are functioning as general acid and base residues in the mechanism of phosphodiester hydrolysis (48). The proposed catalytic mechanism of GlpQ involves the anchimeric assistance of the C2 hydroxyl group, thus requiring this group to be unmodified (i.e. not glycosylated or alanylated at the C2 hydroxyl group of GroP) (41).

**Figure 2.**
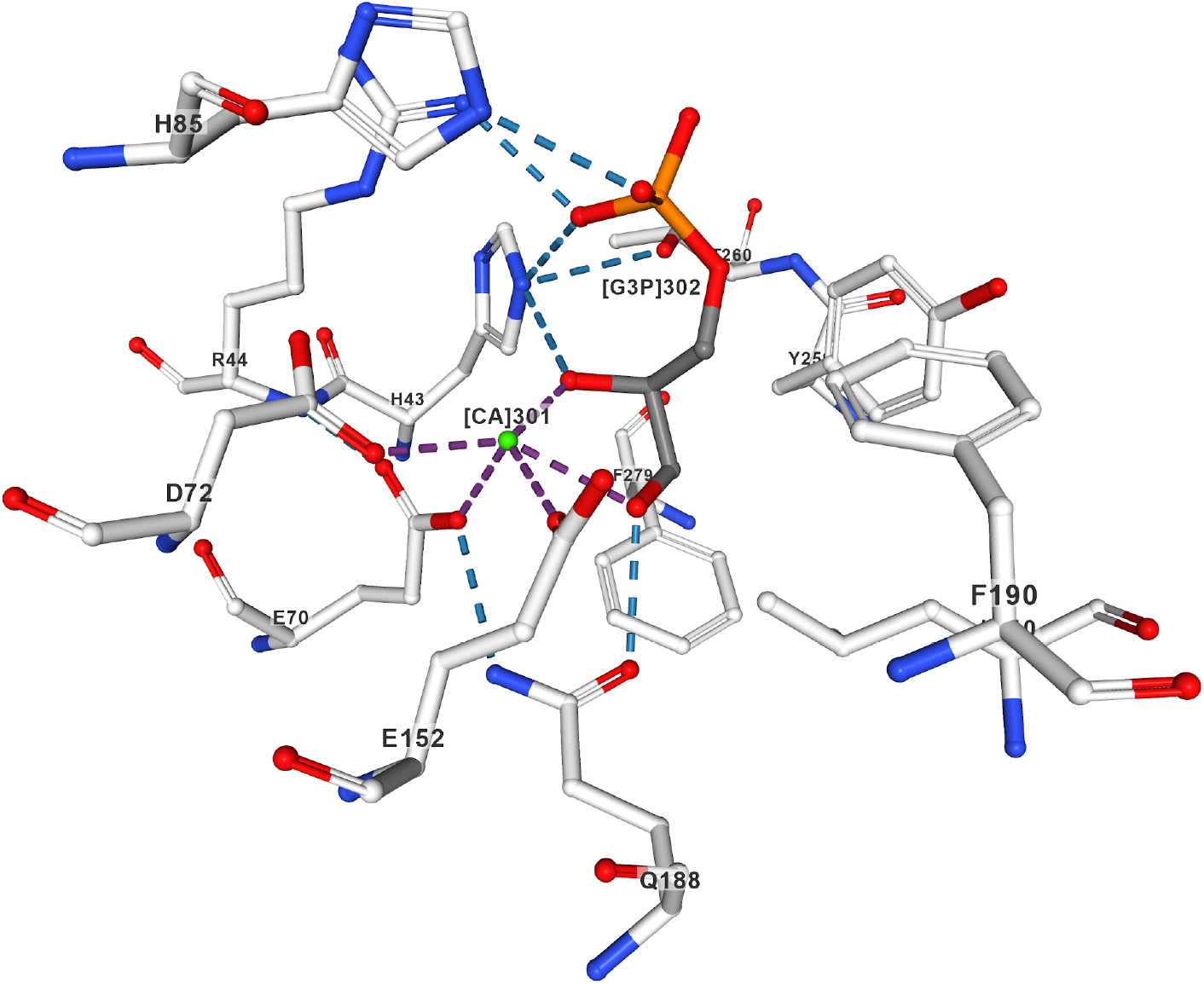
The co-crystal structure of GlpQ with bound Gro3P (pdb identifier: 5T9B; (41)) rationalizes the strict stereospecificity of GlpQ for *sn*-glycero-3-phosphoryl groups. The active site of GlpQ features a hydrophilic side with His85, His43, Arg44, Asp72, Gln188, Glu152 and Glu70 (left of Gro3P; color code: carbon chains grey; oxygens, red; phosphor, orange, nitrogens, blue). The other side of the binding cleft (right of Gro3P) consists of hydrophobic amino acids like phenylalanine, tyrosine and leucine (Phe190, Tyr259, Phe279). The Ca^2+^ion adopts a pentagonal bipyramidal coordination and is coordinated by Glu70, Glu152, and Asp72) as well as by the two hydroxyl groups of Gro3P. The phosphate of the Gro3P substrate makes hydrogen bond interactions with Arg44, His43 and His85. The C2 hydroxyl group of Gro3P interacts with Ca^2+^and His 43 (these interaction would not be possible with Gro1P). Further, the C3 hydroxyl group of the substrate binds to Ca^2+^and Gln188. The structure rationalizes GlpQ’s specificity for *sn*-glycero-3-phosphoryl groups. The C2 hydroxyl group of Gro1P would phase the hydrophobic side and no interaction with Ca^2+^and His 43 would be possible.

### GlpQ sequentially cleaves unmodified WTA by an exo-lytic mechanism

The glycerophosphodiesterase GlpQ of *B. subtilis* has been identified recently as a teichoicase that preferentially digests polyGroP-type WTA lacking modifications on the glycerol subunits (41). However, in this study, product formation with GlpQ had not been followed using polymeric teichoic acids as substrates and thus, neither the strict specificity for unmodified WTA nor the exo-lytic mechanism have been unequivocally shown. We thus aimed at directly monitoring product release by GlpQ from cell wall (PGN-WTA complex) preparations using high performance liquid chromatography-mass spectrometry (HPLC-MS). We first applied cell wall preparations containing glycosylated WTA, which were extracted from *B. subtilis* 168 wild-type cells, and cell wall preparations containing non-glycosylated WTA, which were extracted from ∆*tagE∷erm* cells lacking the WTA alpha-glucosyl transferase TagE (cf. Fig. 1). These samples were digested with GlpQ and product formation was followed by HPLC-MS; Figure 3 depicts the base peak chromatograms (BPC), indicating the total ions in the sample preparations, and the extracted ion chromatograms (EIC) in positive ion mode with (M+H)^+^ = 173.022 m/z, which is the exact mass for GroP, shown in blue. In both cell wall samples, either extracted from wild-type or ∆*tagE∷erm* cells, no GroP was detected in the absence of GlpQ, while upon incubation with the enzyme GroP is released (Fig. 3). Whereas GlpQ releases large amounts of GroP from cell wall samples from ∆*tagE∷erm* cells containing unmodified WTA, and only very little amounts of GroP from glycosylated WTA extracted from wild-type cells (Fig. 3). The amounts of GroP, determined by calculating the area under the curves (AUC), were about 22 times higher when applying cell walls containing unglycosylated WTA (AUC = 5.9 × 10^6^) compared to cell walls containing glycosylated WTA prepared from wild-type cells (AUC = 2.7 × 10^5^), which is in agreement with the proposed chain length of the WTA polymers of 30 - 50 polyol-phosphate repeats. The identity of the GroP reaction product was confirmed by MS and the spectra revealed typical adduct pattern and isotope profiles for GroP (Supporting Information, Fig. S2*A*). It should be noted that with the HPLC-MS method, it is not possible to distinguish between the two stereoisomers of GroP (cf. Fig. 1C). However, given the strict stereospecificity of GlpQ, the product of WTA cleavage has to be *sn*-glycerol-3-phosphate. Very little GroP is released from cell walls containing glycosylated WTA (compare Fig. 3C and D, thus we suggest that GlpQ releases small amounts of non-glycosylated GroP from the free ends of the substrate and stops when encountering a glycosylated (or alanylated) GroP in the chain polymer. Only GroP without modification is released from WTA with GlpQ and no other possible products, neither GroP-Glc, GroP-Ala and other glycosylated or alanylated products, nor larger polymeric products could be detected by HPLC-MS. Thus, GlpQ can be classified as a teichoicase that specifically hydrolyses unmodified *sn*-glycero-3-phosphoryl-WTA.

**Figure 3.**
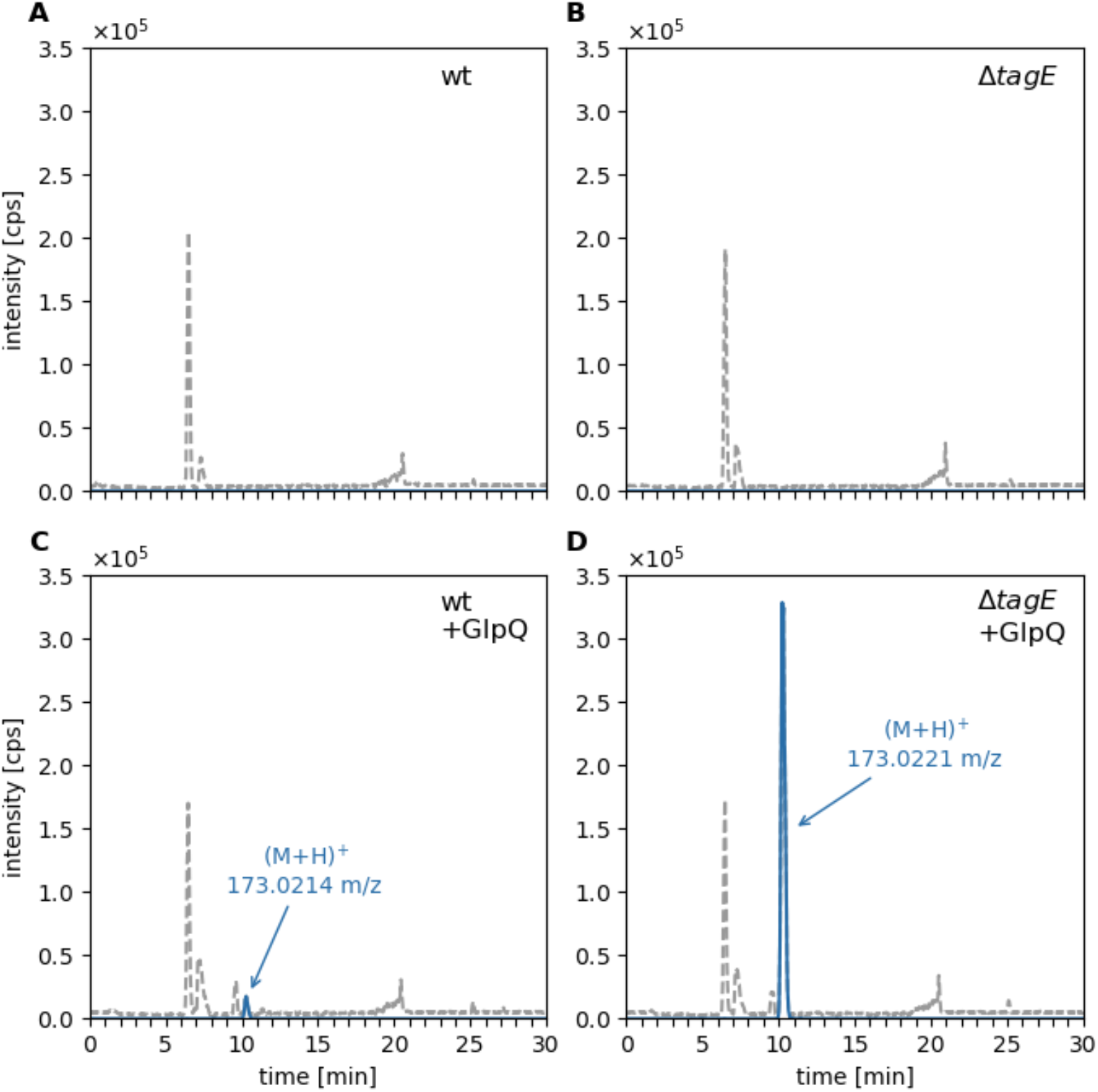
GlpQ releases Gro3P from cell walls of *B. subtilis* **168, predominantly from ∆***tagE* mutant and only little from wild-type (wt) cells. Purified cell wall of *B. subtilis* (containing peptidoglycan and covalently bound WTA) was incubated with GlpQ and the formation of reaction products was analysed by LC-MS. Shown are the base peak chromatograms (BPC) mass range (M+H)^+^ = 120 – 800 (gray dashed) and the extracted ion chromatograms (EIC) of glycerol-phosphate (M+H)^+^ m/z = 173.022 +/− 0.02 (blue solid). ****A****, analysis of wt cell walls (containing partially glycosylated WTA) and ****B****, analysis of ∆*tagE* cell walls without GlpQ added (control). ****C****, analysis of the processing of wt cell walls after incubation with GlpQ for 30 min. The peak area (area under the curve; AUC) of released GroP was AUC = 2.7 × 10^5^. *D*, analysis of the processing of ∆*tagE* cell walls (containing non-glycosylated WTA) digested with GlpQ for 30 min. The obtained AUC = 5.9 × 10^6^ was 22 times as much compared to the release of GroP from wt WTA. No glycosylated or alanylated GroP products were detected.

D-Ala substitutions were removed in the teichoic acid samples by pretreatment as well as applying the GlpQ reaction at pH 8. It has been reported earlier that alanyl esters are rather labile at pH >7, with a half time of hydrolysis at pH 8 and 37°C of 3.9 h (32,50). Accordingly, no difference in the release of GroP was observed, when non-treated and pH 8-pretreated the WTA samples were compared (data not shown). Furthermore, in a time course experiment we observed that already after a few seconds the majority of the product of wild-type and unglycosylated (from *∆tagE* cells) substrate is released by GlpQ (Supporting Information, Fig. S3). Moreover, the amount of GroP released from from unglycosylated substrate did not increase over time (over 2 h of incubation) and remained 22-fold higher than the product released from the wild-type substrate. This indicates that GlpQ has only exo- and no endo-lytic activity and stops when a glycosylated (or alanylated) GroP appears at the free end of the polymer, protecting the rest of the chain from further digestion.

A complete digest of WTA by GlpQ should remove all GroP residues up to the linker disaccharide ManNAc-GlcNAc. To show that this indeed occurs, cell wall preparations (PGN-WTA complex) were thoroughly digested by GlpQ. As the enzyme is rather unstable, GlpQ was added repeatedly: after each round of enzymatic digest for 10 min at 30°C, the supernatant was checked for GroP release by HPLC-MS and then new GlpQ enzyme was added until only very minor additional amounts of GroP were detected. These exhaustively digested cell wall samples were then treated with 5% TCA for 2h at 60°C, whereby the glycosidic phosphodiester bond connecting the WTA linker with the PGN was cleaved. The release of the linker disaccharide was analyzed by HPLC-MS after neutralization of the sample. The identity of the linker disaccharide was confirmed by a mass spectrum revealing typical fragmentations (loss of water), sodium and potassium ion adducts and ^13^C-isotope pattern (Supporting Information, Fig. S2*B*). As control, a complete chemical digest of the PGN-WTA complex was achieved by treatment with 0.5 M NaOH for 2h at 60°C in order to completely remove the GroP chain polymer. Subsequently, the linker disaccharide was released from the latter samples by TCA treatment and analyzed by HPLC-MS. The linker disaccharide was obtained from both wild-type and unglycosylated PGN-WTA complexes by chemical digestion in equal amounts as shown in Figure 4A and B. The amount of linker disaccharide released by chemical digestion was set as to 100% of linker disaccharide in the substrate. As another control, the PGN-WTA complex was treated with TCA alone to determine the amounts of linker disaccharide that TCA can release without requiring NaOH pretreatment. Very small amounts of linker disaccharide (ca. 3.6% of the total) were released from both PGN-WTA variants (Fig. 4C and D). The difference, however, becomes significant once the substrate was pre-digested with GlpQ. While from the wild-type PGN-WTA no more linker was released with GlpQ than with TCA treatment alone, GlpQ was able to digest about 60% of the WTAs up to the linker in the cell wall sample derived from *tagE* mutant cells (Fig. 4E and F).

**Figure 4.**
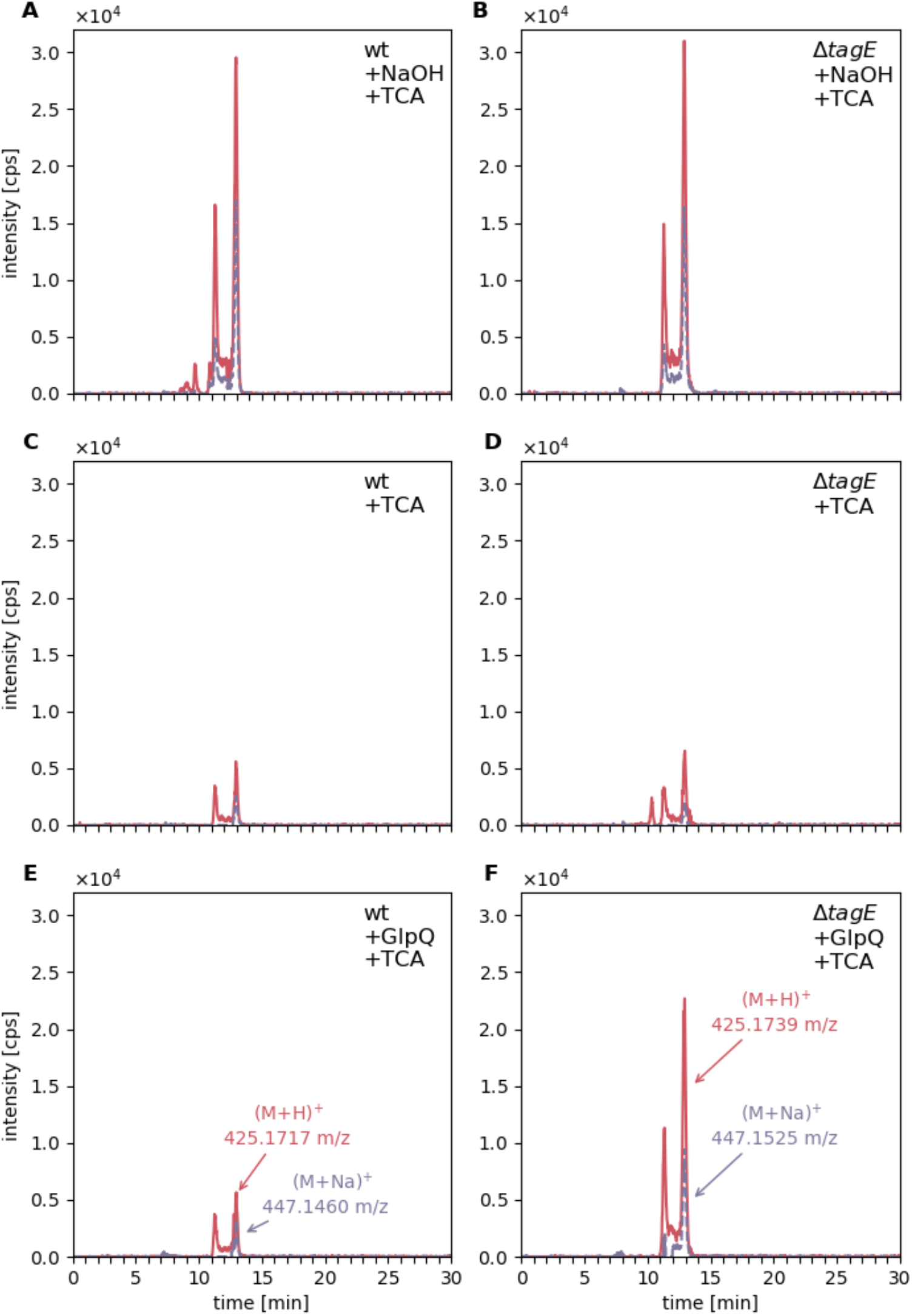
GlpQ completely digests unglycosylated WTA up to the linker disaccharide ManNAc-GlcNAc. Purified cell wall (PGN-WTA complex; 0.1 mg each) of *B. subtilis* 168 wild-type (wt) and ∆*tagE* was repeatedly incubated with GlpQ (seven times for 10 min at 30°C) following treatment of the remaining substrate with 5 % TCA for 2h at 60°C, cleaving the phosphodiester linkage between PGN and the linker disaccharide. The release of ManNAc-GlcNAc was analysed by LC-MS. As a control the cell wall was treated with 0.5 M NaOH for 2h at 60°C to release all GroP from the WTA chain polymers, followed by TCA treatment to release the linker disaccharide. Shown are the extracted ion chromatograms (EIC) of ManNAc-GlcNAc (M+H)^+^ m/z = 425.177 +/− 0.02 (red solid) and (M+Na)^+^ m/z = 447.159 +/− 0.02 (purple dashed). ****A**** and ****B****, complete release of the linker disaccharide after NaOH and TCA treatment from wild-type cell wall (AUC = 15.1 × 10^5^) and from ∆*tagE* cell walls containing unglycosylated WTA (AUC =14.8 × 10^5^). ****C**** and ****D****, linker disaccharide released by TCA treatment alone from wild-type cell wall (AUC= 2.38 × 10^5^) and ∆*tagE* cell wall (AUC = 3.2 × 10^5^). ****E**** and ****F****, linker disaccharide released by TCA treatment after predigest with GlpQ from wild-type cell wall (AUC = 2.84 × 10^5^, i.e. 3.6 % of total linker disaccharide) and ∆*tagE* cell wall (AUC = 1.0 × 10^6^, i.e. 59 % of the totally present linker disaccharide).

### GlpQ cleaves unmodified LTA only after pre-digestion

Since GlpQ specifically cleaves unglycosylated WTA, we next wanted to investigate if also unglycosylated LTA can act as a substrate of the enzyme. Recently the glycosyltransferase CsbB has been shown to be required for glycosylation of LTA in *B. subtilis* (38). LTA was purified from *B. subtilis* wild-type and *∆csbB∷kan* cells using established protocols (51,52). Since LTA has been reported to be extensively modified by D-alanyl esters, we aimed to remove also these modifications prior to GlpQ treatment. In a previous study, it has been shown that incubation of LTA at pH 8.5 for 24 h at room temperature leads to an almost complete removal of D-alanyl esters (22). However, according to these data, also partial degradation occurs during this treatment, albeit the degradation apparently is very limited (the degree of polymerisation dropped from 48 to 43). Since we wanted to avoid absolutely any degradation of the LTA samples, we decided to apply slightly milder conditions, and thus preincubated the LTA preparations in borate buffer at pH 8 for 24 h. The removal of alanine modifications was controlled by NMR (Supporting Information, Fig. S4). ^1^H-NMR spectra of LTA showed characteristic resonances corresponding to D-alanyl ester modifications: signal at δ = 5.35, 4.20 and 1.64 ppm and 4.2 ppm could be assigned to resonances of Gro-2-CH (D-Ala), D-Ala-βH and D-Ala-αH, respectively. These resonances decreased significantly and shifted, indicating a release of D-Ala from the GroP polymer. According to the NMR results, about 70% of the D-alanyl esters were removed by treatment of *B. subtilis* 168 LTA in borate buffer at pH 8 for 24 h at room temperature.

Only very small amounts of GroP were released by GlpQ from LTA extracted from wild-type or *∆csbB* mutant cells, AUC of 1.0 × 10^5^ and AUC of 1.8 × 10^5^ were obtained, respectively (Fig. 5). The experiment was repeated without preincubation under mild alkaline conditions (borate buffer, pH 8, 24 h), which didn’t change the amount of GroP released by GlpQ (Supporting Information, Fig. S3). When incubating the LTA samples under alkaline conditions (0.1 M NaOH, 60°C, 30 min) partial hydrolysis of phosphodiester bonds within the polymer should generate LTA fragments, yet no GroP could be detected, indicating very limited degradation (Figs. 5E and F). Subsequent incubation with GlpQ, however, released substantial amounts of GroP from these LTA preparations, particularly from unglycosylated LTA preparations (Figs. 5G and H). The amount of GroP released by GlpQ was about 3.7-times higher with unglycosylated LTA (AUC = 1.69 × 10^6^) compared to the wild-type LTA (AUC = 4.6 × 10^5^). The same pattern could be observed for LTA obtained from *L. monocytogenes.* While GlpQ released only small amounts of GroP from wild-type (AUC = 6 × 10^4^) and unglycosylated (∆*gtlB*) (AUC = 1.2 × 10^5^) LTA (Figs. 6C and D), the amount increased significantly after NaOH pre-treatment (Figs. 6G and H) with about 4.2-times as much from unglycosylated LTA (AUC = 1.87 × 10^6^) compared to the wild-type (AUC = 4.5 × 10^5^). When LTA was pre-treated with NaOH, the LTA polymer was partially cleaved generating smaller fragments. These fragments possess free ends that expose *sn*-glycero-3-phosphoryl groups (Fig. 7). Subsequent digestion of these fragments with GlpQ released significant amounts of GroP. As seen for digestion of WTA by GlpQ, the enzyme was also releasing more GroP from non-glycosylated LTA fragments, as compared to the glycosylated LTA fragments (see Figs. 5 and 6, G compared to H). These results indicate that GlpQ is only able to release significant amounts of GroP from LTA, when the polymer is pre-cleaved with NaOH, generating LTA-fragments. We can only speculate where the low amounts of GroP released by GlpQ from LTA preparation come from. One possibility is that treatment of LTA preparations at pH 8.0 may cause partial phosphodiester cleavage. However, because phosphodiester cleavage under these conditions is unlikely and analysis of the 1H-NMR spectra of LTA disproves this possibility. Rather, during purification of LTA on a hydrophobic interaction column, lipid II-bound WTA precursors could be co-purified. Since these WTA precursors provide free *sn*-glycero-3-phosphoryl ends the low amount of GroP product released from LTA might come from its degradation.

**Figure 5.**
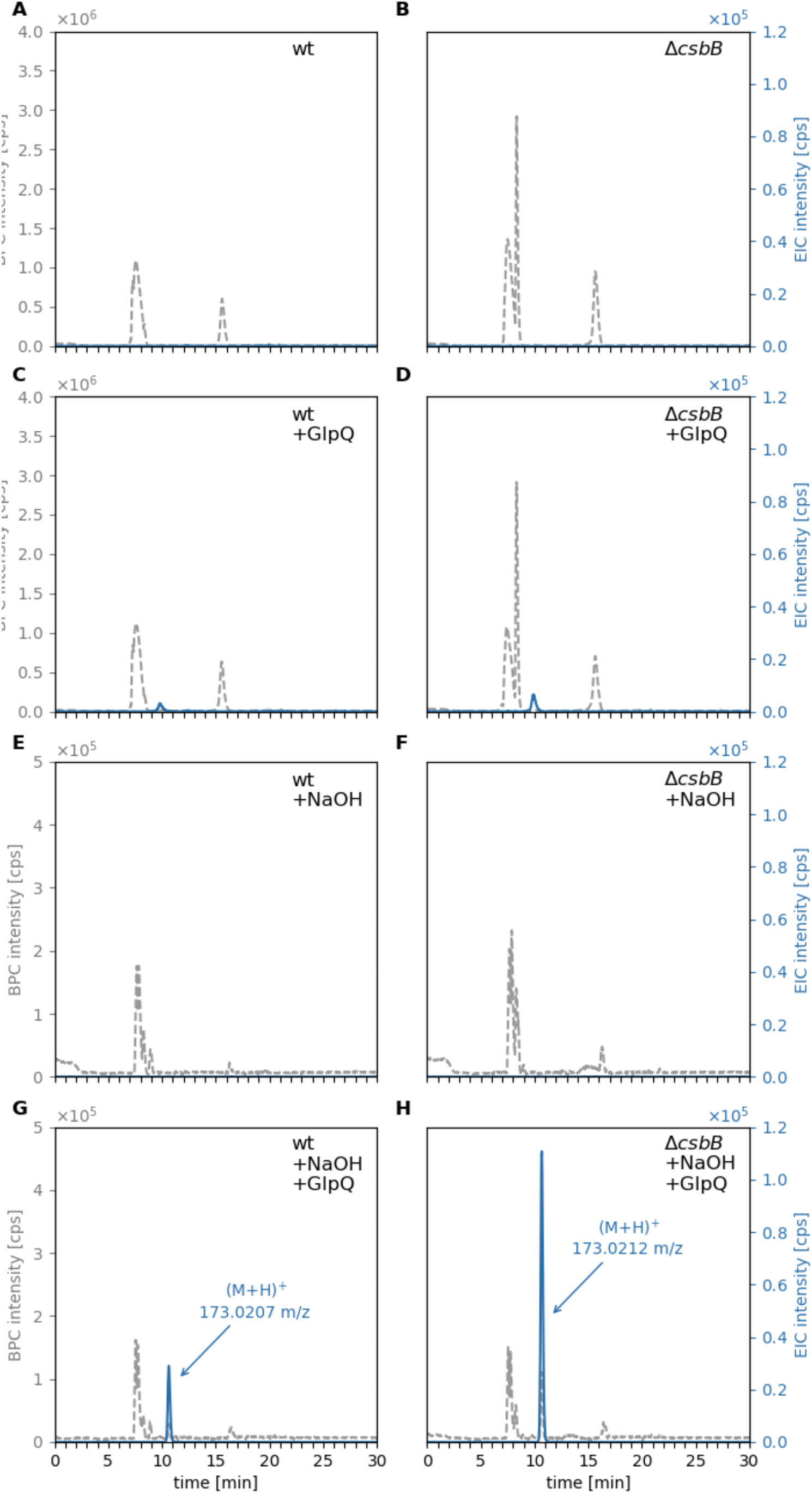
GlpQ releases *sn*-glycerol-3P from NaOH pre-treated LTA of *B. subtilis* wild-type and ∆ *csbB* mutant cells. Purified *B. subtilis* LTA was incubated for 24 h at RT in pH 8 followed by incubated with GlpQ. The formation of reaction products was analysed by LC-MS. Shown are the base peak chromatograms (BPC) mass range (M+H)^+^ = 120 – 800 (gray dashed) and the extracted ion chromatograms (EIC) of glycerol-phosphate (M+H)^+^ m/z = 173.022 +/− 0.02 (blue solid). *A* and *C*, wild-type (wt) LTA (= partially glycosylated LTA) incubated without GlpQ (control) with GlpQ. The peak area of released GroP was AUC = 1 × 10^5^. *B* and *D*, non-glycosylated ∆*csbB* LTA incubated without GlpQ (control) and with GlpQ. The peak area of released GroP was AUC = 1.8 × 10^5^. *E* and *G* wild-type (wt) LTA pre-treated with NaOH incubated without GlpQ (+NaOH) and with GlpQ. The peak area of released GroP was AUC = 4.6 × 10^5^. *F* and *H*, non-glycosylated ∆*csbB* LTA pre-treated with NaOH incubated without GlpQ (+NaOH) and with GlpQ. The peak area of released GroP was AUC = 1.69 × 10^6^.

**Figure 6.**
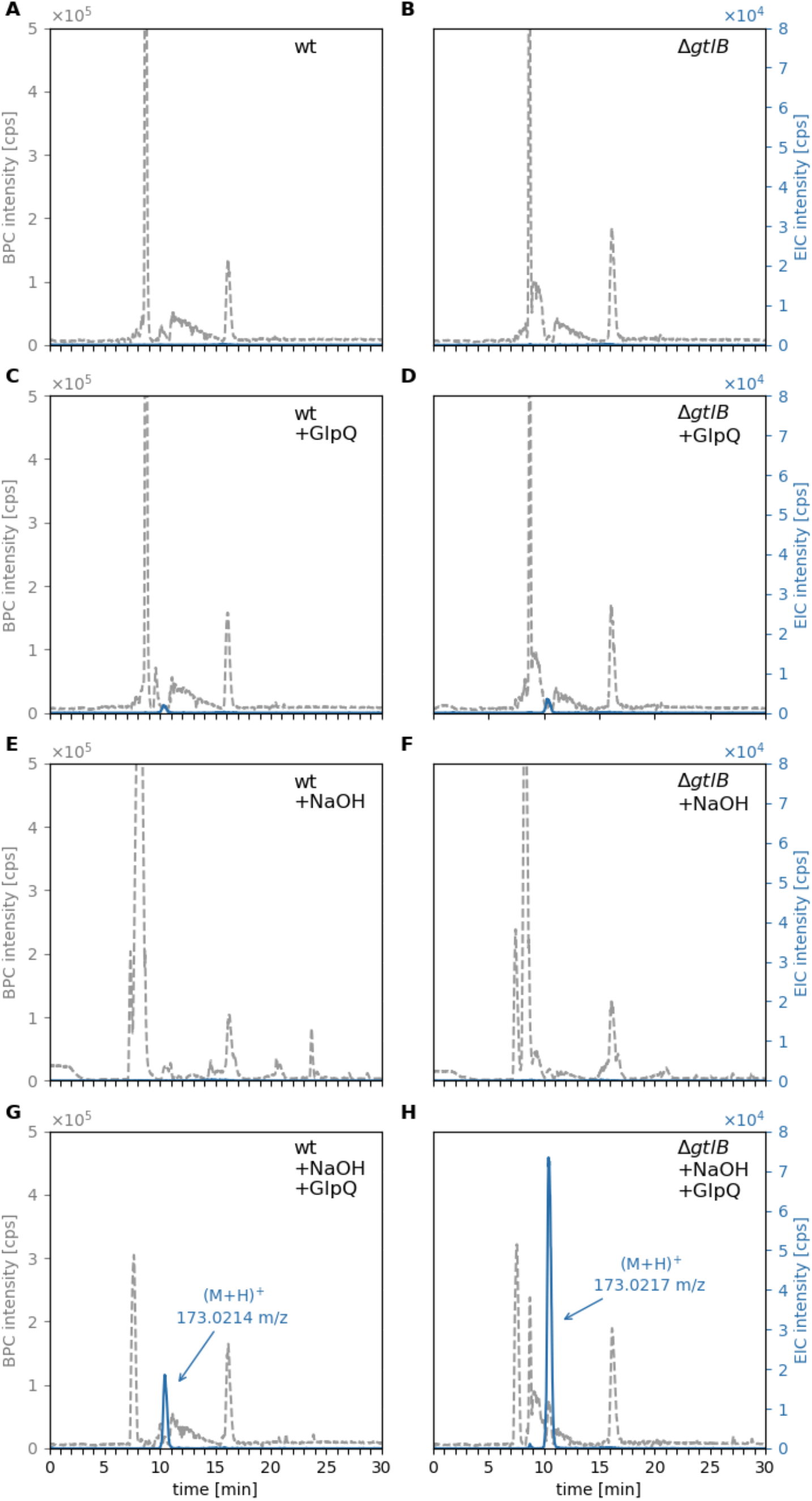
GlpQ releases *sn*-glycerol-3P from NaOH pre-treated LTA of *L. monocytogenes* wild-type and ∆ *gtlB* mutant cells. Purified *L. monocytogenes* LTA was incubated with GlpQ and the formation of reaction products was analysed by LC-MS. Shown are the base peak chromatograms (BPC) mass range (M+H)^+^ = 120 – 800 (gray dashed) and the extracted ion chromatograms (EIC) of glycerol-phosphate (M+H)^+^ m/z = 173.022 +/− 0.02 (blue solid). *A* and *C*, wild-type (wt) LTA (= partially glycosylated LTA) incubated without GlpQ (control) with GlpQ. The peak area of released GroP was AUC = 6 × 10^4^. *B* and *D*, non-glycosylated ∆*gtlB* LTA incubated without GlpQ (control) and with GlpQ. The peak area of released GroP was AUC = 1.2 × 10^5^. *E* and *G*, wild-type (wt) LTA pre-treated with NaOH incubated without GlpQ (+NaOH) and with GlpQ. The peak area of released GroP was AUC = 4.5 × 10^5^. *F* and *H*, non-glycosylated ∆*gtlB* LTA pre-treated with NaOH incubated without GlpQ (+NaOH) and with GlpQ. The peak area of released GroP was AUC = 1.87 × 10^6^.

**Figure 7.**
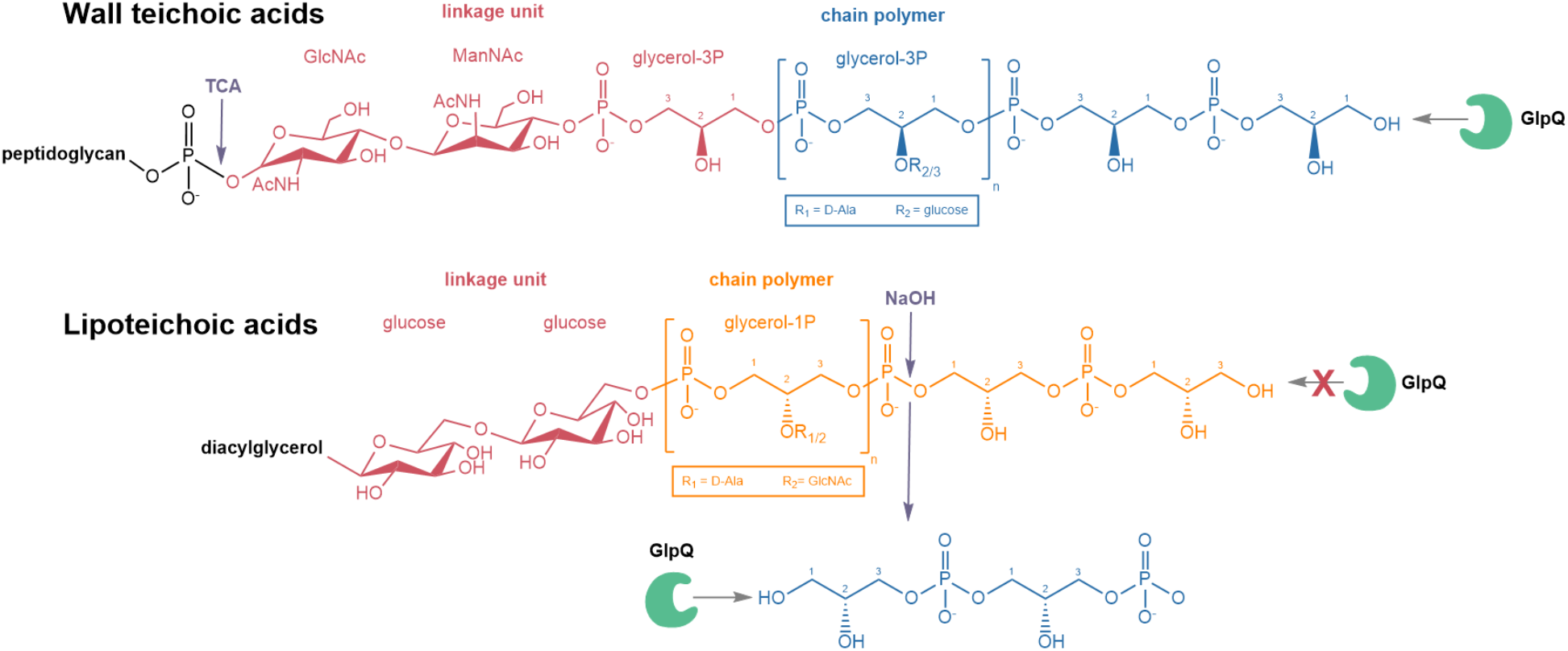
Differential digestion of WTA and LTA with stereospecific *sn*-glycerol-3P phosphodiesterase GlpQ. WTA of *B. subtilis* 168 is phosphodiester polymer made of Gro-3P subunits, that are generally substituted at the hydroxyl group at the C2 position to a certain degree with D-alanine (R_1_) or alpha-glucose (R_2_). Via a linkage unit (red), consisting of a disaccharide, ManNAc-β(1-4)-GlcNAc, and an unmodified Gro-3P, the WTAs are linked to the C6 of MurNAc of the peptidoglycan via a phosphodiester bond. Using trichloro acetic acid (TCA) the glycosidic phosphodiester bond connecting the WTA linker with the PGN can be cleaved. LTA of *B. subtilis* 168 is a phosphodiester polymer made of Gro-1P subunits, hence represents an enantiomer of the WTA polymer. It can be modified at C2 position with D-alanine or GlcNAc and linked to a diacylglycerol via a glucose-β-1,6-glucose linker disaccharide. GlpQ is able to cleave off Gro-3P from the terminal ends of WTA. Conversely, GlpQ is not able to chip off Gro-1P from the terminal ends of LTA. The orientation of the hydroxyl group on the C2 is distinguished by the stereospecific enzyme GlpQ. Treatment with NaOH allows to pre-cleave phosphodiester bonds within the LTA chain polymer, resulting in fragments that contain Gro-3P terminal ends. From these ends GlpQ is able to cleave off Gro-3P moieties. The differential cleavage of WTA and LTA by GlpQ reveals different stereochemistry of the polymers.

In summary, GlpQ releases GroP in significant amounts from WTA (see Fig. 3), but only very small amounts from LTA preparations (see Fig. 5C and 6C). These findings confirm differences in the stereochemistry of the polyglycerol-phosphate polymers WTA and LTA experimentally. In agreement with the described stereospecificity of GlpQ, WTA consists of Gro3P and LTA consist of Gro1P repeating units. The origin for this difference can be found already in the early stages of biosynthesis. WTA biosynthesis starts with Gro3P, which is transferred to CTP by TagD with the simultaneous release of PP_i_, generating CDP-glycerol. The chain polymer is elongated in the cytoplasm with the addition of Gro3P to the growing chain and CMP is released (see Fig. 1A) (53). In contrast, LTA biosynthesis starts with phosphatidylglycerol-CMP, onto which PgsA transfers Gro3P while releasing CMP. The 3P group of Gro3P is released and the product PG is translocated across the membrane and polymerized. The glycerolphosphate group of PG carries a 1-phosphate group, and thus Gro1P is added to the growing chains LTA (see Fig. 1B) (10).

## CONCLUSION

This work reveals the distinct stereoisomerism of the glycerophosphate polymers WTA and LTA of *B. subtilis* by differential digestion with stereospecific phosphodiesterase GlpQ. Firstly, we showed that the stereospecific *sn*-glycerol-3P phosphodiesterase GlpQ is an exo-lytic hydrolase that sequentially cleaves off GroP entities from WTA, that lack any modification in form of D-alanylation or α-glucosylation, up to the linker unit that connects WTA with the PGN. Secondly, GlpQ is unable to cleave intact, unmodified LTA. Thus, WTA and LTA polymers of *B. subtilis* 168 constitute enantiomers, consisting of Gro3P (WTA) and Gro1P (LTA) building blocks, respectively. Accordingly, limited hydrolysis of LTA with NaOH, which leads to a random cleavage of phosphodiester bonds within the polymer, yields fragments that contain Gro3P terminal ends, from which GlpQ is able to cleave off Gro3P entities. The difference in stereochemistry between the WTA and LTA has critical consequences regarding differential physiological functions, regulation, and turnover of both polymers. The results of this study rationalize the specific interaction of WTA and LTA by stereospecific enzymes and protection against the simultaneous degradation with possibly fatal effects for cell viability.

## Experimental Procedures

### Bacterial Strains and Growth Conditions

The bacterial strains, plasmids and oligonucleotides used in this study are listed in supplemental Table S1. *Bacillus subtilis* 168 wild-type and ∆*tagE∷erm* strains were obtained from the *Bacillus* genetic stock center (Columbus, Ohio, USA). *B. subtilis* ∆*csbB∷kan*, *Listeria monocytogenes* wild-type strain 10403S and ∆*gtlB∷strep* mutant were obtained from the Gründling lab (38). These bacteria were used for the isolation of whole cell wall (peptidoglycan-WTA complex) and teichoic acid preparations. They were cultured at 37°C in lysogeny broth (LB broth Lennox, Carl Roth) with continuous shaking at 140 rpm or on solid LB supplemented with 1.5% agar. Overnight cultures (~16 h) were used to inoculate fresh LB medium and grown to yield an optical density at 600 nm (OD_600_) of 1. Cells were harvested by centrifugation (3000 × *g*, 20 min, 4°C). *E. coli* BL21 (DE3) cells (New England Biolabs) were used to heterologously express recombinant GlpQ phosphodiesterase from *B. subtilis*. These cells, transformed with pET28a-*glpQ*, were grown in LB medium supplemented with 50 μg/ml kanamycin until OD_600_ 0.7 was reached, followed by induction with 1 mM isopropyl β-D-thiogalactopyranoside and further propagation for 3 h. Cells were harvested by centrifugation (3000 × *g*, 20 min, 4°C) and used for the purification of recombinant GlpQ.

### Construction of plasmids and purification of recombinant GlpQ

*B. subtilis* 168 *glpQ* was amplified by PCR with the primers pET28a-glpQ-for and pET28a-glpQ-rv (MWG Eurofins, Ebersberg, Germany). Oligonucleotide primers are listed in supplemental Table S1. The PCR products were purified (Gene JET purification kit and Gene Ruler, 1-kb marker, Thermo Fisher Scientific) and then digested with appropriate restriction enzymes (New England Biolabs) and ligated with T4 DNA ligase (Thermo Fisher Scientific) into the expression vector pET28a (Novagen), allowing to overproduce a C-terminal His^6^-tag fusion protein. *E. coli* BL21 (DE3) cells carrying pET28a-*glpQ* were grown as described above and lysed in a French pressure cell. The His-tagged GlpQ protein was purified by Ni^2+^-affinity chromatography using a 1 ml HisTrap column (GE Healthcare) followed by size exclusion chromatography on a HiLoad 16/60 Superdex 200 pg column (GE Healthcare) and purity was checked with a 12% SDS-PAGE. The purity of the enzyme was confirmed via SDS-PAGE (see figure 2 A). From a 1 l culture 3.6 mg GlpQ have been obtained. The enzyme was stored with a concentration of 0.23 mg/ml at −20°C in 0.1 M Tris-HCl buffer (pH 8).

### Biochemical Characterization of GlpQ

To determine the enzymatic properties of GlpQ, 1 pmol of pure recombinant enzyme was incubated with 10 mM GPC. The reaction was stopped by adding 200 μl of pH 3.3 buffer (0.1% formic acid, 0.05% ammonium formate) and the released glycerol-phosphate was measured by HPLC-MS. For the pH stability, GlpQ was pre incubated in buffers at different pHs (pH 2: Clarks and lubs, pH 3-6: acetic acid, pH 6-7 MES, pH 7-9: Tris, pH 10: NaHCO3) for 30 min at 30°C before adding 5 μl each to a 45 μl mix with 0.1 M Tris pH 8 buffer and substrate for 5 min. The pH optimum was tested by incubating GlpQ with 10 mM GPC for 5 min in buffers with different pH. For the temperature stability GlpQ was pre incubated in 0.1 M Tris-HCl, pH 8 at different temperatures ranging from 4 to 75°C for 30 min, followed by a 5 min incubation with GPC at 30°C and pH 8.0. The optimum temperature was tested by incubating GlpQ for 5 min at different temperatures with GPC at pH 8.

### Preparation of Cell Wall, WTA and LTA

For the preparation of cell walls (peptidoglycan-WTA complex) 2 L of *B. subtilis* 168 wild-type or ∆*tagE∷erm* cultures (exponential growth phase, OD_600_ = 0.9) were harvested and resuspended in 30 ml piperazine-acetate buffer (50 mM, pH 6) with 12 U proteinase K and boiled for 1 h. The cytosolic fractions were removed by centrifugation (3000 × g, 15 min, 4°C). The pellet was resuspended in 6 ml buffer (10 mM Tris, 10 mM NaCl, 320 mM imidazole, adjusted to pH 7.0 with HCl) and 600 μg α-amylase, 250 U RNase A, 120 U DNase I and 50 mM MgSO_4_ were added. The sample was incubated at 37°C for 2 h while shaking, 12 U Proteinase K was added the incubation continued for 1 h. 4% SDS solution was added 1:1 and the mixture was boiled for 1 h. The SDS was removed by repeated ultracentrifugation steps (20 times at 140 000 × g, 30 min, 40°C) and suspension in H_2_O_bidestilled_ as well as dialysis against H_2_O_bidestilled._ The SDS content was controlled with the methylene blue assay described earlier (54). The cell wall preparation was dried in a vacuum concentrator.

LTA from *B. subtilis* 168 (wild-type and ∆*csbB*) and *L. monocytogenes 10403S* (wild-type and ∆*gtlB*) was prepared by butanol extraction and purification by hydrophobic interaction chromatography using a 24 × 1.6-cm octyl-Sepharose column, according to published protocols (51,52).

### Teichoic Acid Digestion with GlpQ and Analysis of Glycerol-phosphate Release

WTA assays were conducted in 0.1 M Tris-HCl buffer (pH 8, supplemented with 1 mM CaCl_2_) with 0.1 mg cell wall preparation (peptidoglycan with attached WTA from *B. subtilis* 168 wild-type and ∆*tagE∷erm)* as a substrate and 0.7 μM GlpQ. The samples were incubated for 30 min at 30°C.LTA assays occurred in 0.1 M Tris-HCl buffer (pH 8, supplemented with 1 mM CaCl_2_) with 0.2 mg LTA extract (*B. subtilis* 168 wild-type and ∆*csbB∷erm)* and 0.7 μM GlpQ in a total volume of 50 μl. The samples were incubated for 1 h at 30°C. LTA was pre-digested by incubating with 0.1 M NaOH for 30 min at 60°C, followed by neutralization with HCl and drying in the vacuum concentrator.

Sample analysis was conducted using an electrospray ionization-time of flight (ESI-TOF) mass spectrometer (MicrOTOF II; Bruker Daltonics), operated in positive ion-mode that was connected to an UltiMate 3000 high performance liquid chromatography (HPLC) system (Dionex). For HPLC-MS analysis 7 μl of the sample supernatant was injected into a Gemini C18 column (150 by 4.6 mm, 5 μm, 110 Å, Phenomenex). A 45 min program at a flow rate of 0.2 ml/min was used to separate the compounds as previously described (55). The mass spectra of the investigated samples were presented as base peak chromatograms (BPC) and extracted ion chromatograms (EIC) in DataAnalysis program and were presented by generating diagrams using Python 3.6 with the Matplotlib (version 2.2.2) library.

## ACKNOWLEDGMENTS

C.M., J.R, and A. P. acknowledge funding by the Deutsche Forschungsgemeinschaft (DFG, German Research Foundation). C.M. receives DFG grants SFB766, Project-ID 398967434- TRR 261 and Project-ID 174858087- GRK1708, J.R. revieves DFG grant RI 2920/1-1 and A.P. receives DFG grants SFB766, TRR 34 and TRR 156. This work was further supported by the DFG-funded Cluster of Excellence EXC 2124 Controlling Microbes to Flight Infections. A.G. acknowledges funding from the Wellcome Trust grant 210671/Z/18/Z and MRC grant MR/P011071/1. The authors declare that they have no conflicts of interest with the contents of this article.

---

**Figure S1.**
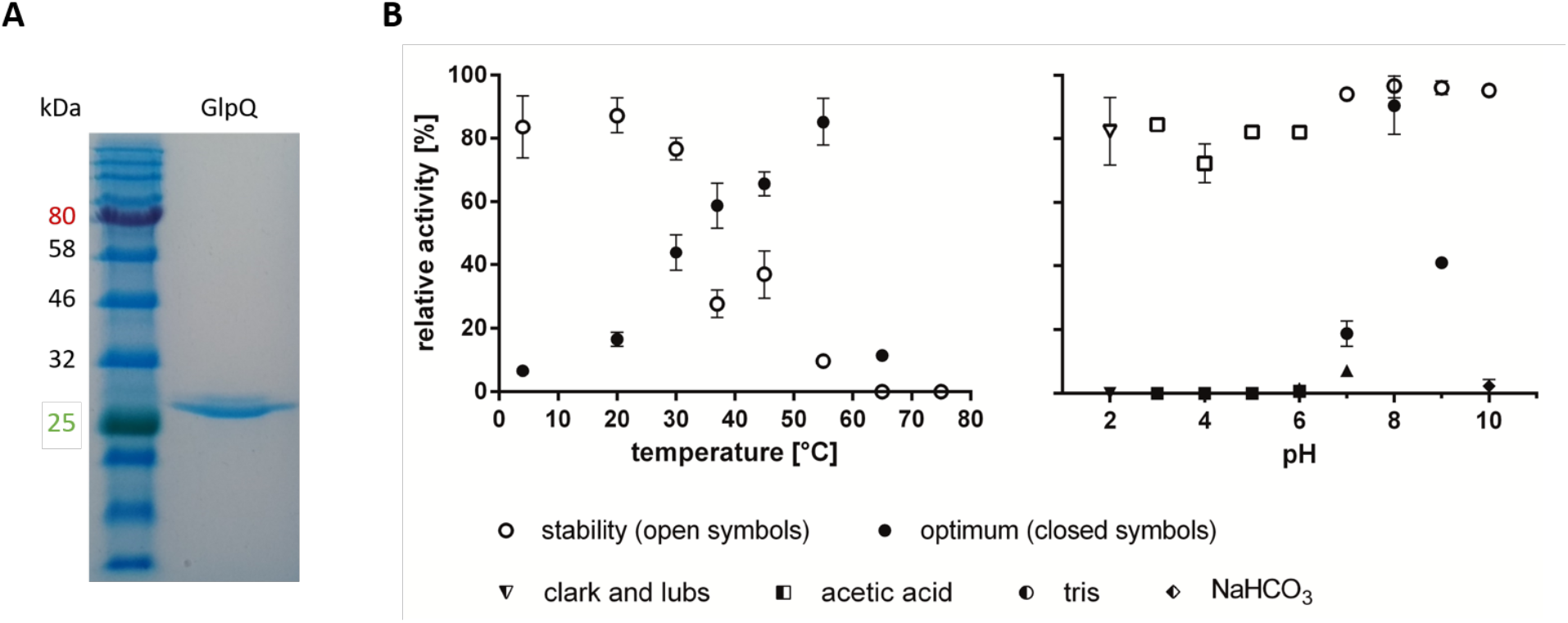
Enzyme purity, stability and optima of recombinant GlpQ. ***A*** shows the purity of heterologously expressed recombinant GlpQ-His_6_fusion protein as analysed by SDS-PAGE after purification of the enzyme by Ni^2+^affinity chromatography and size exclusion chromatography (1 ug protein was loaded on a 12% polyacrylamide gel). A single band is visible just above 25 kDa size marker, in agreement with the calculated molecular weight of GlpQ-His^6^without signal peptide of 29.6 kDa. ****B****, shows the temperature and pH characteristics of GlpQ. The enzyme is stable for 30 min at temperatures up to 30°C, but stability rapidly drops at temperatures above 30°C within this time frame. Activity of GlpQ increased with temperature up to 55°C and reveals only about half maximum activity at 30°C. GlpQ is stabile within a broad range between pH 2-10 in the indicated buffers and has a very sharp pH-optimum at 8.0. In all assays, 1 pmol GlpQ was incubated with 10 mM GPC and reaction product was analyzed by LC-MS, after 30 min of incubation at 30°C. For temperature stability and optimum, 100% relative activity reflect area under the curve (AUC)-values of 2240 and 4088, respectively. For pH stability and pH optimum 100% AUC were 28571 and 31903, respectively.

**Figure S2.**
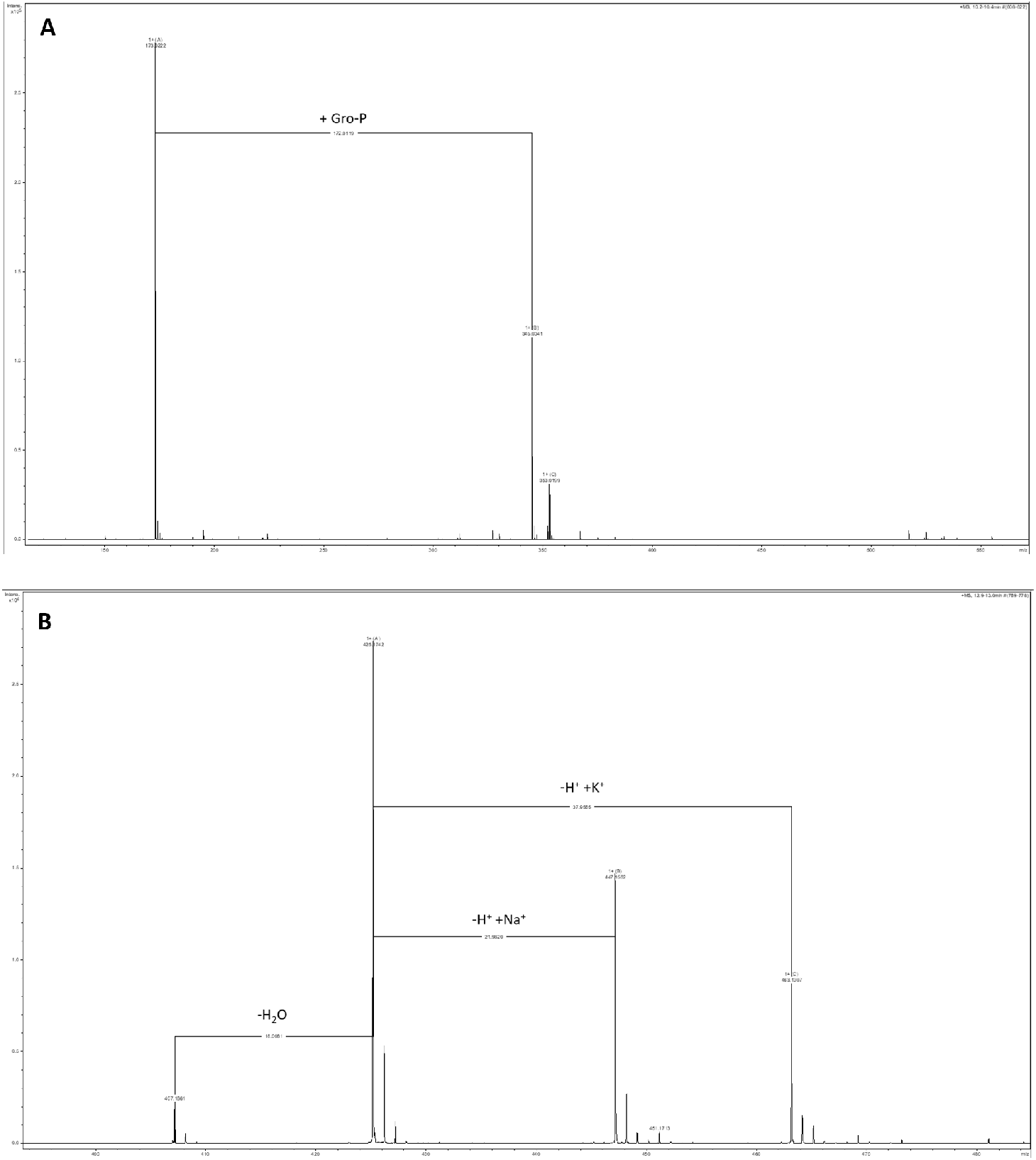
Mass spectra of glycerol-phosphate (GroP) and the WTA linker disaccharide ManNAc-β-1,4-GlcNAc. ***A***, shows the mass spectrum of GroP detected in positive ion mode [M+H]^+^ (experimental 173.0222; theoretical monoisotopic mass 173.0210), also revealing a non-covalently bound GroP dimer, [2M+H]^+^ (experimental 345.0341; theoretical 345.0346). ****B****, shows the mass spectrum of the WTA linker disaccharide, ManNAc-GlcNAc, detected in positive ion mode [M+H]^+^(experimental 425.1742; theoretical monoisotopic mass 425.1766), also revealing the sodium adduct [M+Na]^+^ (experimental 447.1562; theoretical 447.1585), the potassium adduct [M+K]^+^ (experimental 345.0341; theoretical 345.0346), asl well as a product of neutral water loss, [M-(H_2_0)+H]^+^ (experimental 407.1661; theoretical 407.1660).

**Figure S3.**
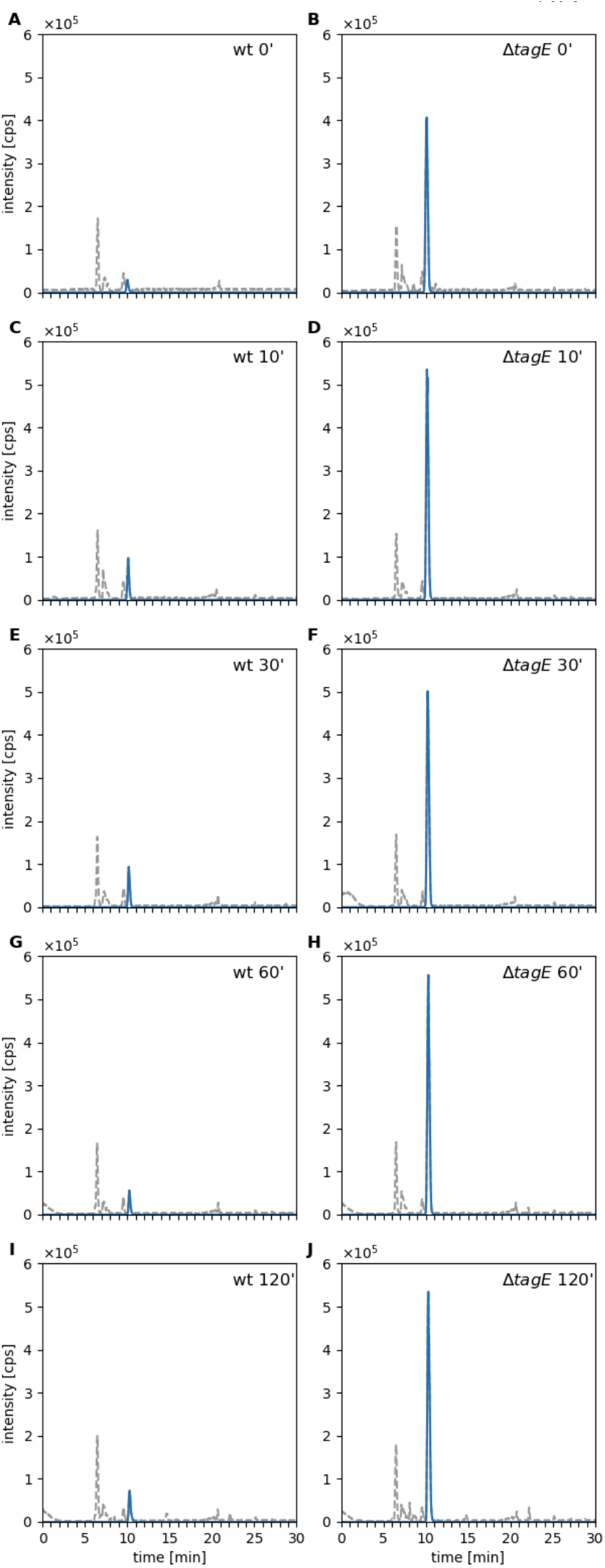
Time course of WTA digest by GlpQ. The vast amount of product (GroP; blue lines) is released by GlpQ within the first seconds of incubation of cell walls purified from wild-type cells (wt; ***A***,**C**,**E**,**G**,**I**; incubation time in min) and unglycosylated cell wall (from *∆tagE* cells; ***B***,**D**,**F**,**H**,**J**; incubation time in min). GlpQ releases significantly more product from unglycosylated substrate (from *∆tagE* cells) than from the wild-type. Even over a long period of time on more GroP was released from wild-type cell wall, indicating that GlpQ has only exo- and no endo-lytic activity. 0.25 mg purified cell wall of *B. subtilis* (containing PGN and covalently bound WTA) was incubated with 0.7 μM GlpQ and the formation of reaction products was analysed by LC-MS. Shown are the base peak chromatograms (BPC) mass range (M+H)^+^ = 120 – 800 (gray dashed) and the extracted ion chromatograms (EIC) of glycerol-phosphate (M+H)^+^ m/z = 173.022 +/− 784 0.02 (blue solid). The reaction was stopped by incubation at 95°C followed by LC-MS analysis.

**Figure S4.**
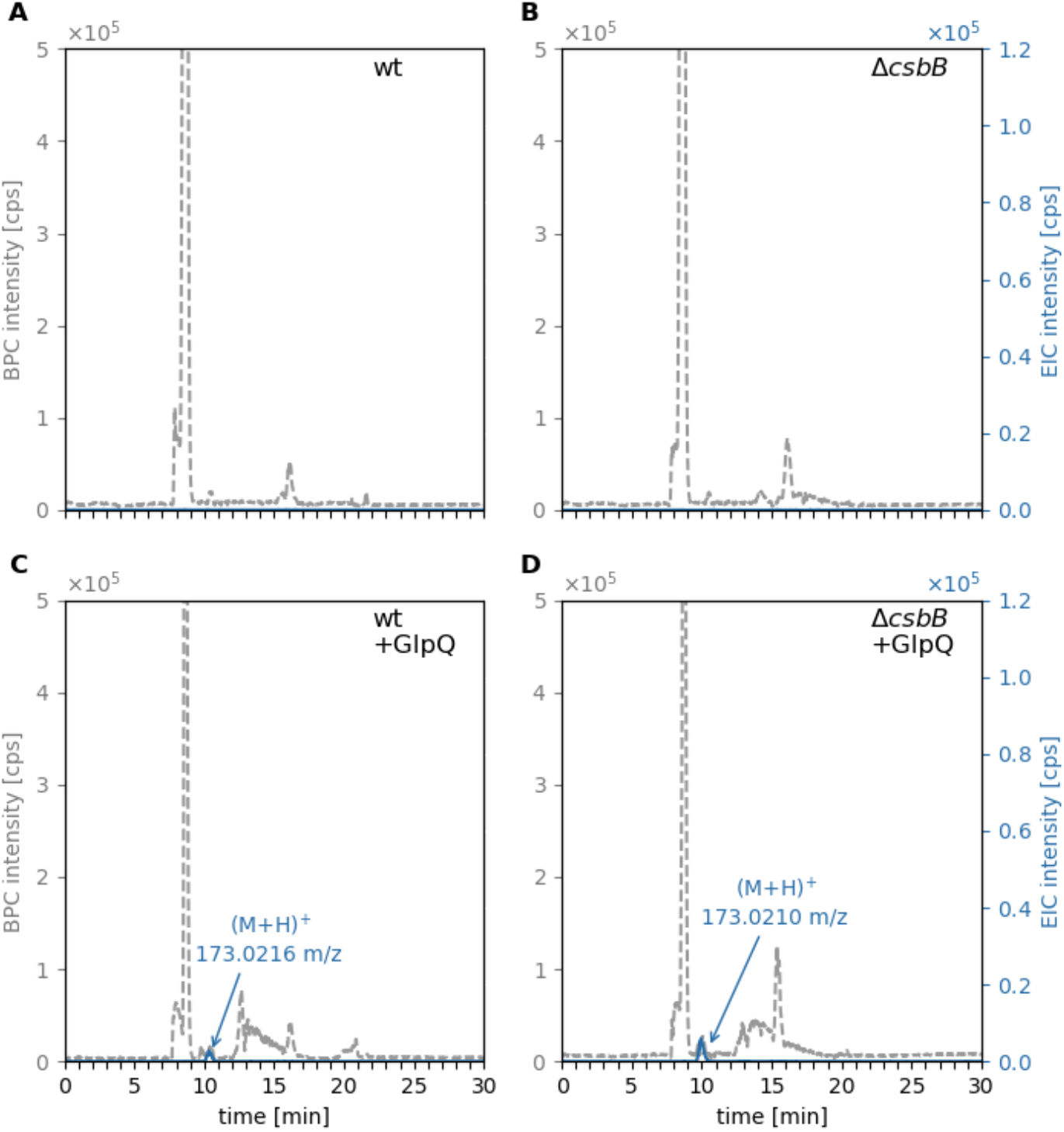
*B. subtilis* 168 LTA not preincubated at pH 8 cannot be cleaved by GlpQ. Purified *B. subtilis* LTA was incubated with GlpQ and the formation of reaction products was analysed by LC-MS. Very little amounts of GroP were released by GlpQ. ****A**** and ****C****, wild-type (wt) LTA (= partially glycosylated LTA) incubated without GlpQ (control) with GlpQ. The peak area of released GroP was AUC = 6 × 10^4^. ****B**** and ****D****, non-glycosylated ∆*csbB* LTA incubated without GlpQ (control) and with GlpQ. The peak area of relased GroP was AUC = 1.4 × 10^5^. Shown are the base peak chromatograms (BPC) mass range (M+H)^+^= 120 – 800 (gray dashed) and the extracted ion chromatograms (EIC) of glycerol-phosphate (M+H)^+^ m/z= 173.022 +/− 0.02 (blue solid).

**Figure S5.**
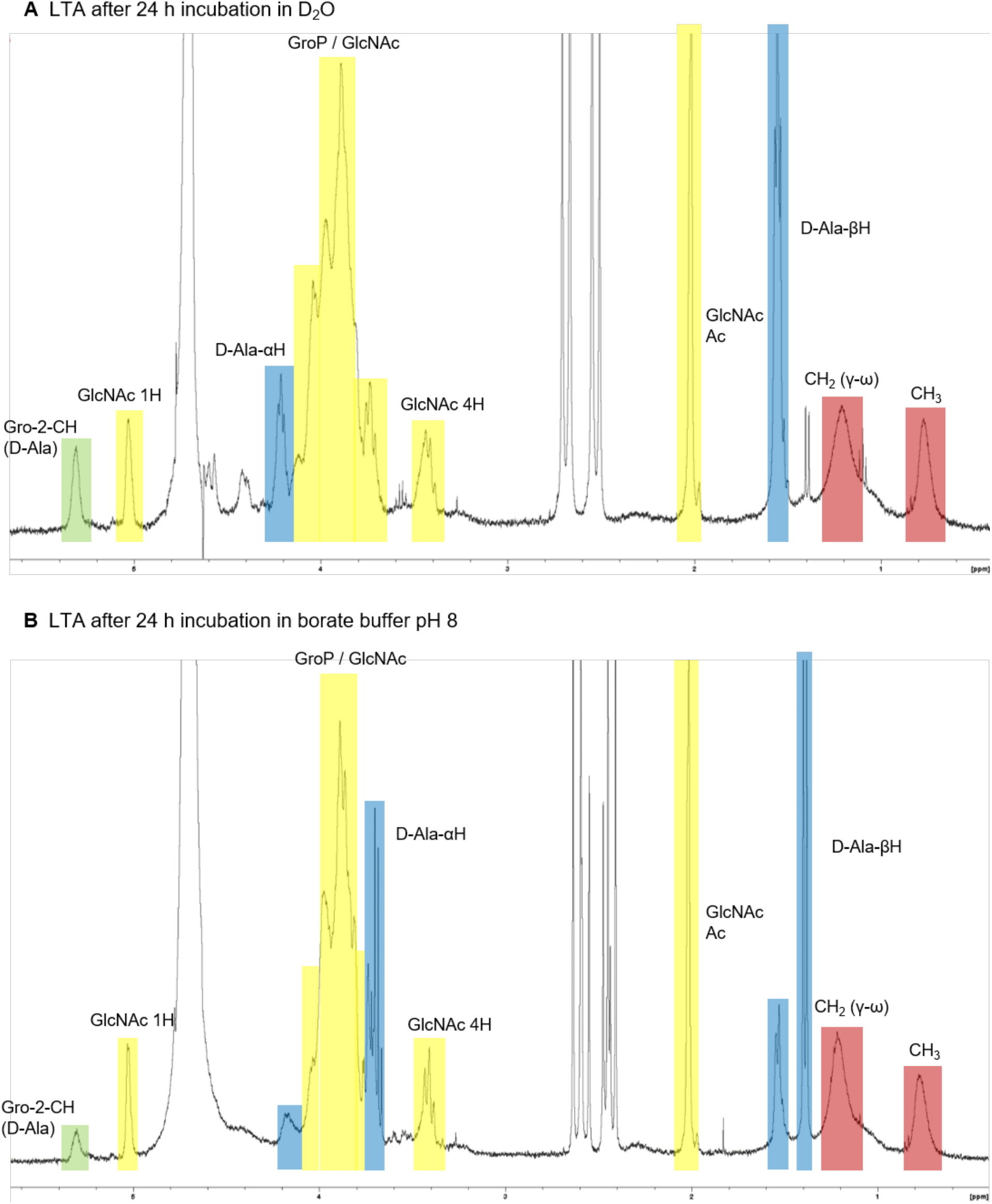
^1^H-NMR analysis reveals severely reduced D-alanyl ester modifications when LTA was incubated for 24 h at pH 8. Shown are the ^**1**^H-NMR spectra (400 MHz, 303K) of LTA isolated from *B. subtilis* 168 wt (2 mg) either incubated for 24 h at RT in D_2_O, pH 7.0 *A* or in D_2_O containing 0.1 M borate buffer pH 8.0 *B*. Color coding identifies signals indicating removal of D-alanyl residue from the GroP polymer (green): the resonance of the methine group of *sn*-glycerol (Gro-2-CH) containing an D-alanyl ester (D-Ala) is reduced and partially shifted from 5.3 ppm to 3.9 ppm and the D-Ala-ɑH and D-Ala-βH resonances (blue) are significantly reduced and partially shifted in the LTA sample incubated at pH 8 compared to LTA in D_2_O. Other resonances assigned to GlcNAc substitution and the GroP polymer (yellow) and to the fatty acids of LTA (red) are not influenced by incubation in borate buffer at pH 8.0. From the signal integrales it was estimated that about two-thirds of the D-Ala substituents were removed in the LTA sample by incubation at pH 8 for 24 h. NMR analysis was performed on a 400-MHz Bruker Advance III spectrometer at 303 K with a TCl cryoprobe. NMR spectra were interpreted according to (22).

**Table S1.**
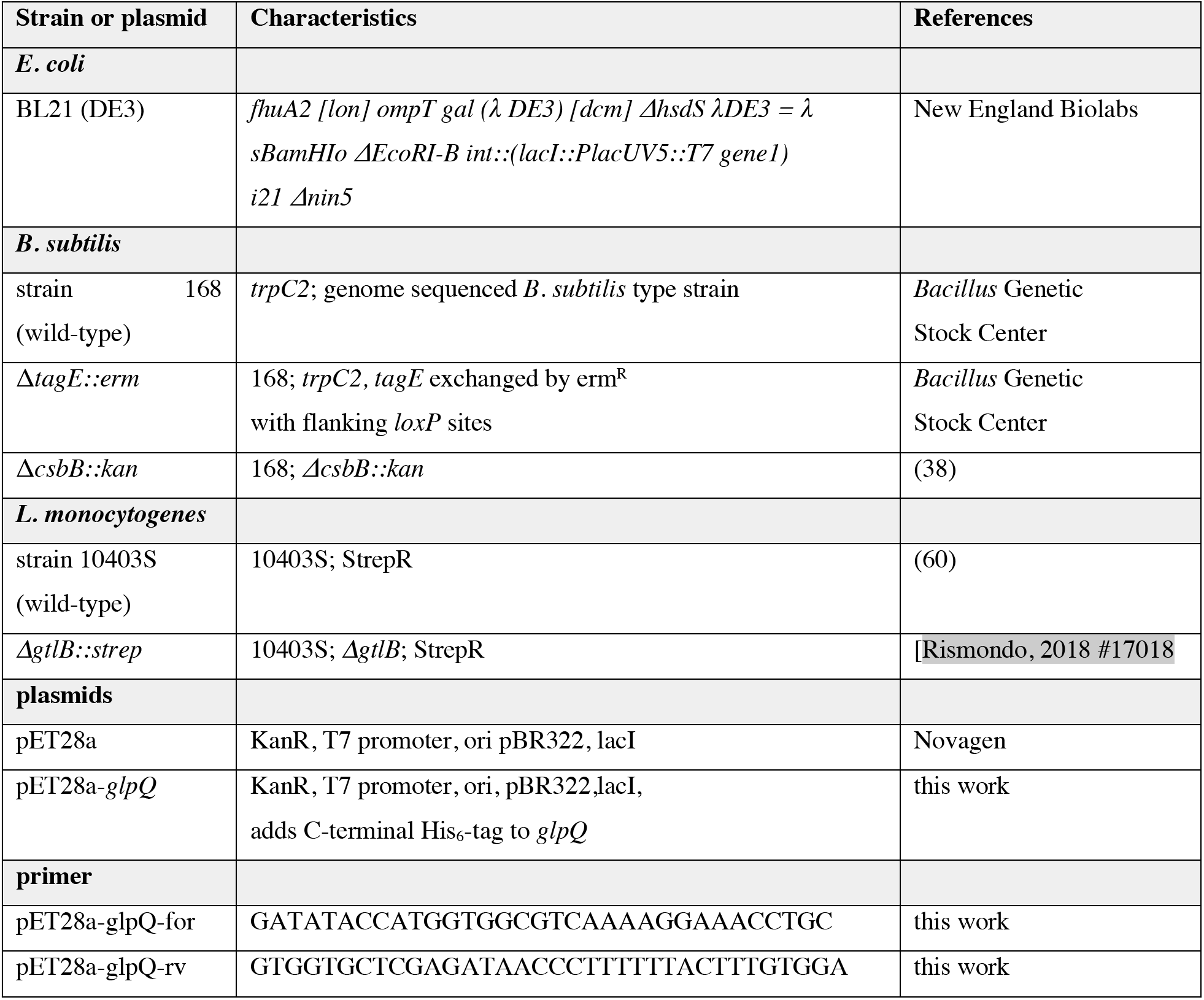
Strains, plasmids and primer used in the study.

